# Alcohol and Vapourized Nicotine Co-Exposure During Adolescence Contribute Differentially to Sex-Specific Behavioural Effects in Adulthood

**DOI:** 10.1101/2021.06.23.449629

**Authors:** Jessica Ruffolo, Jude A. Frie, Hayley H.A. Thorpe, M. Asfandyaar Talhat, Jibran Y. Khokhar

## Abstract

**Introduction:** Co-occurrence of e-cigarette use and alcohol consumption during adolescence is frequent. However, little is known about their long-lasting effects when combined. Here, we examined whether adolescent co-exposure to alcohol drinking and vapourized nicotine would impact reward- and cognition-related behaviours in adult male and female rats during adulthood.

**Methods:** Four groups of male and female Sprague Dawley rats (n=8-11/group/sex) received either nicotine (JUUL 5% nicotine pods) or vehicle vapour daily between postnatal days 30-46, while having continuous voluntary access to ethanol and water during this time in a two-bottle preference design. Upon reaching adulthood, rats underwent behavioural testing utilizing Pavlovian conditioned approach testing, fear conditioning and a two-bottle alcohol preference test.

**Results:** A sex-dependent effect was found in the two-bottle preference test in adulthood such that females had a higher intake and preference for alcohol compared to males regardless of adolescent exposure; both male and female adult rats had greater alcohol preference compared to adolescents. Male rats exposed to vapourized nicotine with or without alcohol drinking during adolescence exhibited altered reward-related learning in adulthood, evidenced by enhanced levels of sign-tracking behaviour. Male rats that drank alcohol with or without nicotine vapour in adolescence showed deficits in associative fear learning and memory as adults. In contrast, these effects were not seen in female rats exposed to alcohol and nicotine vapour during adolescence.

**Conclusions:** The present study provides evidence that co-exposure to alcohol and vapourized nicotine during adolescence in male, but not female, rats produces longterm changes in reward- and cognition-related behaviours.

**Implications:** These findings enhance our understanding of the effects of alcohol drinking and nicotine vapour exposure in adolescence. Moreover, they highlight potential sex differences that exist in the response to alcohol and nicotine vapour, underscoring the need for follow-up studies elucidating the neurobiological mechanisms that drive these sex differences, as well as the long-term effects of alcohol and nicotine vapour use.

## INTRODUCTION

E-cigarettes have become increasingly popular among adolescents^1^; in 2020, 30.7% of 10^th^ graders and 34.5% of 12^th^ graders reported using e-cigarettes, following a two-fold increase over the past two years^1^. An equally important issue is the consumption of alcohol by adolescents. Over 78% of adolescents have consumed alcohol by late adolescence, and 15% meet the criteria for DSM-IV lifetime alcohol abuse^2^. Alcohol and nicotine are frequently used sequentially and simultaneously by adolescents^3^ with a high prevalence of concurrent e-cigarette vaping and alcohol consumption^4^. Vaping high school students also showed greater alcohol drinking compared to non-vapers^5^, and alcohol and e-cigarette use is the most common type of co-use in this population^6^. The long-term effects of vaping are currently unknown, as are the consequences of e-cigarette/alcohol co-use^6^. In contrast, the additive effects of combustible tobacco and alcohol co-use are well established. For example, cigarette smoking amplifies cognitive deficits in adults who excessively drink alcohol, and alcohol-dependent adults who smoke cigarettes show pronounced neuropsychological damage compared to alcohol-dependent non-smokers^7,8^. Unfortunately, the consequences of nicotine and alcohol co-use in adolescence on subsequent behaviour in adulthood are limited.

The adolescent brain undergoes critical neuronal and structural development, making adolescence a period of vulnerability to the effects of drugs^9^, with brain imaging studies suggesting altered brain structure and function in adolescent users compared to non-users^9^. However, the causal consequences of adolescent co-use cannot be systematically examined in humans and thus, animal models are required to explore the cause-effect relationship. Although there is a dearth of pre-clinical studies that have examined the acute effects of adolescent nicotine and alcohol co-exposure, the research that has been conducted suggests their co-use to have additive effects on behavioural outcomes. In adolescent male rats, concurrent intravenous self-administration of nicotine and alcohol was more reinforcing than either drug alone – an effect not observed in adults^10^. A combination of the two substances also increases ambulatory activity, and decreases anxiety-like behaviours in adolescent, but not adult, males^11^. Together, these results suggest that there may be fundamental differences in the effects of alcohol and nicotine co-use in adolescents compared to adults.

Despite the paucity of preclinical research that explores the long-term ramifications of adolescent nicotine and alcohol co-use, both drugs have been examined in isolation. Nicotine or alcohol exposure in adolescence increases the risk of substance use later in life, where alcohol exposure in adolescent rats increased voluntary ethanol drinking and preference in adulthood^12^, and adolescent nicotine-exposed rodents showed increased vulnerability to nicotine’s reinforcing effects and enhanced reward responses to other drugs as adults^13^. These drug-induced neuroadaptations may result in the sensitization of reward-incentive processes such that reward-related stimuli acquire enhanced salience^14^. Repeated exposure to alcohol or nicotine in adolescent rats increased conditioned approach toward reward-associated cues in adulthood^15–17^. Moreover, adult rodents exhibit long-term impairments across several cognitive domains as a consequence of alcohol exposure in adolescence, including spatial working memory^18^, object recognition memory^19^ and fear retention^20^. Similarly, rats treated with nicotine during adolescence display long-lasting dysfunctions in attention, impulsive behaviour^21^ and serial pattern learning^22^.

To date, no preclinical studies have directly tested the effects of adolescent alcohol drinking and nicotine vapour co-exposure on reward- and cognitive-related behaviours in adulthood. Given the known sex differences in the response to alcohol and nicotine^23^, as well as sex differences in brain development^24^, it is imperative to study these effects in both male and female rodents. We hypothesize that the co-exposure to nicotine vapour and alcohol in adolescence would produce pronounced changes in reward- and cognitive-associated behaviour when compared with exposure to either drug alone. Specifically, we predicted that the co-exposure to alcohol and vapourized nicotine would increase alcohol intake and preference in adulthood, enhance the incentive salience of reward-predictive stimuli and impair fear associative learning and memory.

## METHODS

All procedures were approved by the Animal Care Committee at the University of Guelph under Canadian Council on Animal Care Guidelines. Male (N=39) and female (N=35) Sprague-Dawley rats aged post-natal day (PND) 21 upon arrival were obtained from Charles River (Montreal, Canada). The animals were maintained on a 12-hour light-dark cycle (7AM-7PM). Upon weaning (PND23), same-sex littermates were housed two per cage. On PND28, pair-housed rats were separated by a mesh divider in order to measure individual food, water and alcohol intake while still having social contact. All animals had access to standard chow and water *ad libitum* until behavioural testing commenced. The rats were randomly assigned to one of four exposure groups: 1) Vehicle vapour/Water (CO) (N=8/sex); 2) Vehicle/Alcohol (AO) (N=11/9, male/female); 3) Nicotine Vapour/Water (NO) (N=8/sex) or 4) Nicotine Vapour/Alcohol (AN) (N=11/9, male, female).

### Drug Preparation

#### Nicotine

JUUL mint flavoured 5% nicotine e-liquid pods (59mg/ml) (JUUL Labs, Toronto, Canada) or vehicle e-liquid without flavouring was administered. The vehicle e-liquid was a mixture of 30:70 propylene glycol to vegetable glycerine, representative of commercially available JUUL pods^25^.

#### Alcohol

Ethanol (Commercial Alcohols, Brampton, Canada) was diluted to a final concentration of 10% v/v in tap water^12^.

### Vapour Administration Procedure

Beginning at PND30, passive nicotine vapour exposure was conducted using the OpenVape apparatus^26^. Animals in the nicotine vapour only (NO) and concurrent alcohol drinking and nicotine vapour (AN) exposure groups received nicotine vapour. Animals in the control (CO) and alcohol only (AO) exposure groups received vehicle vapour. All animals received 10 minutes of vapour exposure per day for 16 days, with each epoch of pump activation resulting in 10, 2-second puffs/minute^26^. Animals from each exposure group and sex were placed in the exposure chamber together.

### Adolescent Alcohol Exposure

During the duration of vapour exposure (PND30-46), rats assigned to AO and AN exposure groups were given 24-hour access to 10% ethanol and tap water in a two-bottle preference design (described in detail below). Rats assigned to the CO and NO exposure groups received continuous access to two bottles of tap water.

### Two-Bottle Preference Test

The effects of adolescent voluntary alcohol drinking (with or without nicotine vapour inhalation) on adulthood alcohol consumption and preference were quantified using a two-bottle alcohol preference test^27^. Beginning at PND110, all rats were given 24-hour access to 10% ethanol and tap water for four days. To prevent location preference development, the position of the bottles was switched daily. Bottle weights were recorded daily. The main outcome measurements were alcohol consumption (in g/kg) and alcohol preference. Alcohol preference was calculated as:

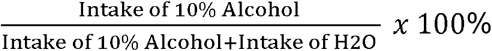

### Effects of Adolescent Nicotine Vapour and Alcohol Co-Exposure an Reward-Related Behaviour an Adulthood

In adulthood, body weights were reduced to 85% of baseline body weight. All rats remained food restricted until the completion of the experiment and were fed after each testing session.

#### Pavlovian Conditioned Approach Apparatus

Pavlovian conditioned approach (PCA) task was used to measure the incentive salience of reward-predictive cues. Behavioural procedures were carried out in standard operant chambers (Coulbourn Instruments, Allentown, PA) enclosed in sound-attenuating cubicles (76.2 × 46.99 × 44.96 cm). The chambers were outfitted with a food cup located in the center of the right sidewall and retractable levers were positioned to the left and right of the food cup. The cubicles also contained surveillance cameras to monitor behaviours during testing. Data collection from Graphic State software (Coulbourn Instruments, Allentown, PA) was transformed using a customized Microsoft Excel macro.

#### Behavioural Procedure

All rats were given 45mg Dustless Precision sucrose banana flavoured pellets (Bio-Serv, Flemington, NJ, product #F0024) in their home cages for one day to reduce potential neophobia. On the first day of the experiment (PND85), each rat was assigned to a chamber and received one 30-minute magazine training session, during which a single banana pellet was freely delivered on a random time 30-second schedule. PCA testing began 24-hours following magazine training and lasted 12 days. Each daily testing session was 60 minutes in duration and consisted of 25 conditioned stimulus (CS)+ and 25 CS- trials with an average inter-trial interval (ITI) of 60 seconds. CS+ trials consisted of a 10-second extension of a lever followed by the delivery of 2 banana pellets upon lever retraction. During CS- trials, a 10-second extension of the other lever occurred, but no reinforcer was delivered upon retraction. The presentation of the CS was pseudorandom such that no more than two of the same CS presentations could occur consecutively. The assignment of the left and right levers as either CS+ or CS-was counterbalanced across animals and within exposure groups.

### Effects of Adolescent Vapourized Nicotine and Alcohol Co-Exposure on Associative Memory in Adulthood

#### Fear Conditioning Apparatus

Experiments were conducted in four standard fear-conditioning chambers (Med Associates Inc., St. Albans, VT; 29.53 × 23.5 × 20.96 cm) enclosed in sound-attenuating cubicles (63.5 × 36.83 × 74.93 cm). A tone generator presented an auditory cue (90dB, 2-kHz) as the CS. The steel bars were wired to a shock generator to deliver an electric foot shock (1.0 mA) as the unconditioned stimulus. Chambers were outfitted with either a lemon or vanilla scent and flat or zig-zag grid floors as contextual cues, with use of these cues counterbalanced across animals within each exposure group. Freezing behaviour, defined as total motor immobility except for movement necessitated by respiration^28^, was recorded and analyzed using Video Freeze Software. On the cue testing day, animals were introduced to a novel context arrangement (e.g., zig-zag grid floor substituted for flat grid floors, lemon substituted for vanilla scent) and a plastic sheet that rounded the chamber walls was inserted to provide a new context.

#### Behavioural Procedure

As previously described^29^, the training session consisted of five 10-second tone presentations followed by foot-shock delivery during the last 2 seconds of the tone with an ITI of 64 seconds. Fear conditioning began on PND103 with the first trial starting 3 minutes after the rat is placed in the chamber. After a 24-hour period, freezing to the context was assessed by returning the rats to their original chamber for 8 minutes, with no tone or shock presented. One day later, fear conditioning to the tone was assessed by placing rats in a novel context. Following an initial 30-second delay, the tone was presented 20 times for 10-seconds each with a 30-second ITI between tone presentations. No shock was delivered to the animals. Incidences of freezing were recorded during the ITIs on training and tone test sessions, and in 64 second bins during the context test session^29^.

### Statistical Analyses

Data analysis was performed using IBM SPSS Statistics 27 (Armonk, New York, United States) software. All data will be available upon publication at https://www.khokharlab.com/.

#### Two-Bottle Preference Test Data Analysis

Average alcohol intake and preference in adolescence and adulthood were compared both within- and between- ages using a 2-way between-subjects ANOVA with adolescent nicotine exposure and age as between-subjects factors, for males and females in groups exposed to alcohol in adolescence. Average alcohol intake and preference data from adulthood only was analyzed by a 2-way between-subject ANOVA with adolescent exposure and sex as between-subjects factors.

#### PCA Behavioural Data Analysis

Behavioural dependent measures included: (1) number of lever presses and food cup entries per session, (2) probability of pressing the lever and entering the food cup during a trial and (3) the latency to press the lever and enter the food cup during the 10-second CS presentation. Given previously observed sex differences in the impact of adolescent nicotine and alcohol exposure on behaviour^22,23^, males and females were analyzed separately for each behavioural measure using a 2-way-repeated measures factorial ANOVA with exposure (nicotine vapour and alcohol) as a between-subjects factor and day as a within-subject factor. Average PCA index scores (described below) for males and females were analyzed with a 2-way ANOVA with exposure (nicotine vapour and alcohol) as the between-subjects factor. Cohen’s d was also calculated as an estimation of effect size. The interpretation criteria for effect sizes were considered small, medium, or large if they corresponded to partial η^2^ and Cohen’s *d* of at least 0.0099 (*d* = 0.29-0.49), 0.0588 (*d* = 0.50-0.79) and 0.13790 (*d* = ≥ 0.80), respectively.

#### Quantification of Sign- and Goal-Tracking Behaviour

A PCA index score was calculated to classify animals as goal-trackers, sign-trackers or intermediates based on average performance during all 12 sessions^30^. The PCA Index was calculated as the sum of (1) Response Bias (contacting the CS+ lever or food cup entries in relation to total number of CS+ or food cup responses), (2) Probability Difference (difference between the probability of pressing the CS+ lever and probability entering the food cup) and (3) Latency Score (the difference between latency to contact the CS+ lever and latency to enter the food cup), which was then divided by 3. Animals with scores of −1.0 to −0.3 were classified as goal-trackers and animals with scores of +0.3 to +1.0 were classified as sign-trackers. Animals that were within the range of - 0.29 to +0.29 were defined as intermediates.

#### Fear Conditioning Data Analysis

Separate 2-way analyses were conducted for males and females given that they display distinct patterns of fear expression^31^. Freezing behaviour was analyzed using repeated measures ANOVA on each of the three days with adolescent exposure (nicotine vapour and alcohol) as the between-subjects factor and trial or block (64-second epoch) as the within-in subjects factor. One male animal in the CO and two male animals in the AN group were excluded due to their freezing being more than 2 standard deviations outside the mean.

## RESULTS

### Two-Bottle Preference Test

#### Female rats consumed more alcohol than males in adulthood, with both sexes showing greater alcohol preference in adulthood compared to adolescence

No main effect of exposure or an exposure by sex interaction was revealed for alcohol intake or preference in adolescence or adulthood. A 2-way ANOVA comparing alcohol preference from adolescence and adulthood revealed a main effect of age for males [F(1,40) = 8.3; p = 0.006, η_p_^2^ = 0.173] and females [F(1,32) = 30.8; p < 0.05, η_p_^2^ = 0.491] where preference for alcohol was higher in adulthood relative to adolescence **(Figure 1 A,B).** A significant main effect of age for average alcohol intake (g/kg) was observed in females [F(1,32) = 9.3; p = 0.021, η_p_^2^ = 0.156] but not in males, indicating alcohol intake was higher in adulthood relative to intake in adolescence for female rats **(Figure 1 C,D).** An overall increase in alcohol intake [F(1,66) = 19.8; p < 0.05, η_p_^2^ = 0.231] and preference [F(1,66) = 21.4; p < 0.05, η_p_^2^ = 0.245] in adulthood was detected in female rats compared to male rats across all exposure groups **(Figure 1 E,F)**.

**Figure 1.**
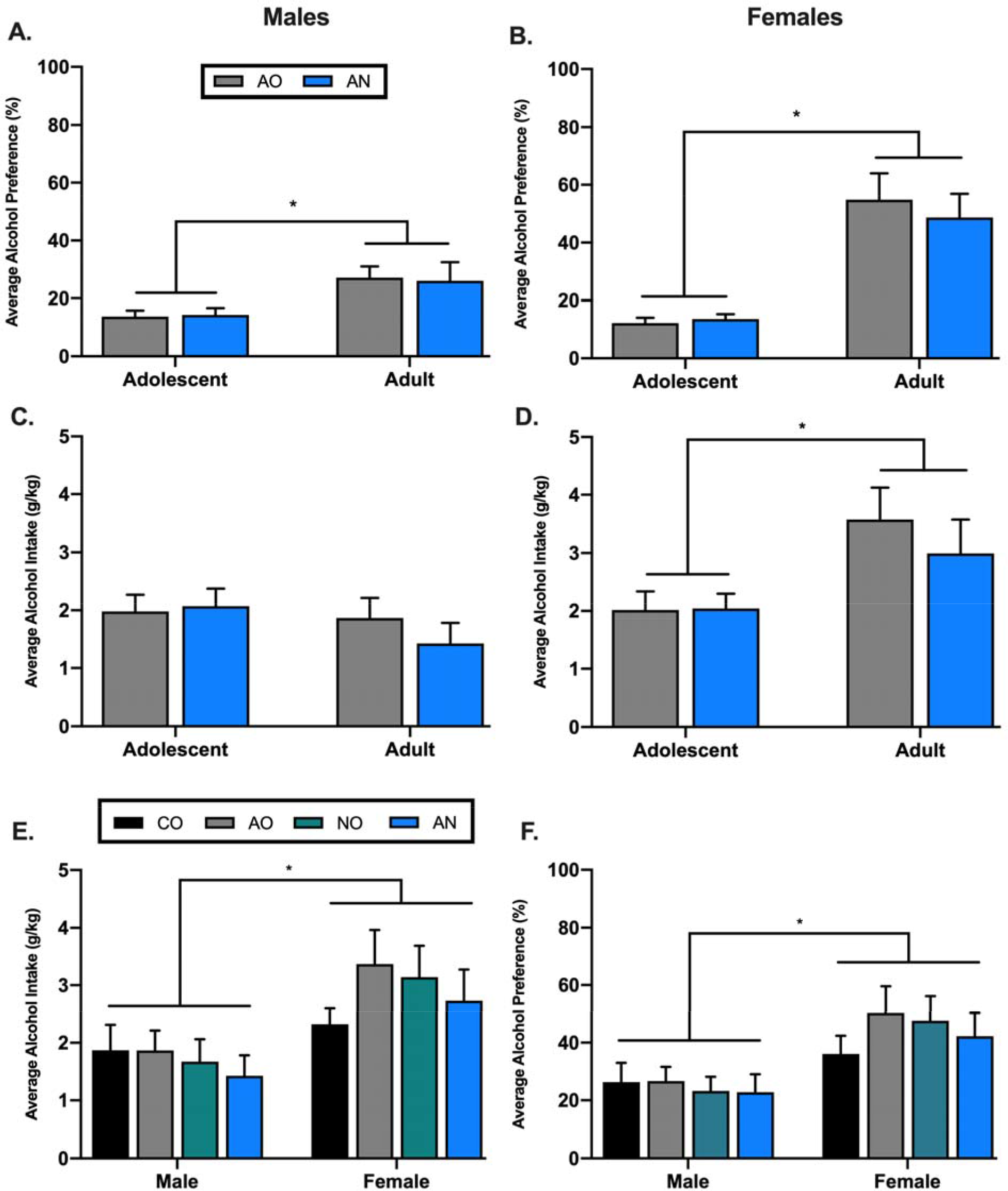
Effects of adolescent nicotine vapour and alcohol exposure on alcohol drinking in adult male and female rats. On average, males A) and females B) in AO and AN exposure groups had a higher alcohol preference during adulthood relative to their preference in adolescence. C) There were no differences in alcohol consumption in males across age. D) Females consumed more alcohol as adults compared to their consumption as adolescents. On average, females E) drank significantly more alcohol in adulthood and F) had a higher preference for alcohol compared to males across all exposure groups. The data is presented as mean ± SEM. * p ≤ 0.05 significant differences between A, B, D) adolescent and adult drinking behaviours and E, F) male and female drinking behaviours.

### Pavlovian Conditioned Approach

#### Male adolescent nicotine vapour exposure altered reward-associated learning in adulthood

Sign-tracking was measured as lever pressing during CS+ presentation over the 12 testing sessions. Analysis of the number of lever presses revealed a main effect of session for males [F(3.3,116.9) = 11.4; p < 0.05, η_p_^2^ = 0.246] and females [F(3.0,94.1) = 9.3; p < 0.05, η_p_^2^ = 0.231] with lever pressing increasing across sessions. A main effect of nicotine vapour was reported for males [F(1,35) = 4.9; p=0.03, η_p_^2^ = 0.123] in the between-subject analysis, with both adolescent nicotine vapour-exposed male groups exhibiting significantly higher lever pressing compared to CO males (*d* = 1.0) **(Figure 2A)**. No effect of nicotine vapour or alcohol drinking was observed in females **(Figure 2B)**. Analysis of the probability to lever press revealed a main effect of session in males [F(3.5,122.9) = 25.6; p < 0.05, η_p_^2^ = 0.422] and females [F(4.0,124.1) = 24.3; p < 0.05, η_p_^2^ = 0.440] with the probability to press the lever increasing across sessions. A main effect of nicotine vapour exposure was revealed in males [F(1,35) = 9.5; p = 0.004, η_p_^2^ = 0.214] where both nicotine vapour exposed groups displayed a higher probability to press the lever compared to CO (*d* = 1.2) and AO (*d* = 0.8) exposure groups **(Figure 2C)**. This effect was not observed in females **(Figure 2D)**. Analysis of latency to lever press revealed a main effect of session in males [F(3.4,117.8) = 24.9; p < 0.05, η_p_^2^ = 0.416] and females [F(4.0,123.4) = 24.7; p < 0.05, η_p_^2^ = 0.443] with latency to press the lever decreasing across sessions. Male rats from both groups exposed to vapourized nicotine in adolescence demonstrated a shorter latency to press the lever [F(1,35) = 9.1; p = 0.005, η_p_^2^ = 0.206] relative to CO (*d* = 1.2) and AO (*d* = 0.8) exposure groups **(Figure 2E)**. No differences were present amongst exposure groups in female rats **(Figure 2F)**.

**Figure 2.**
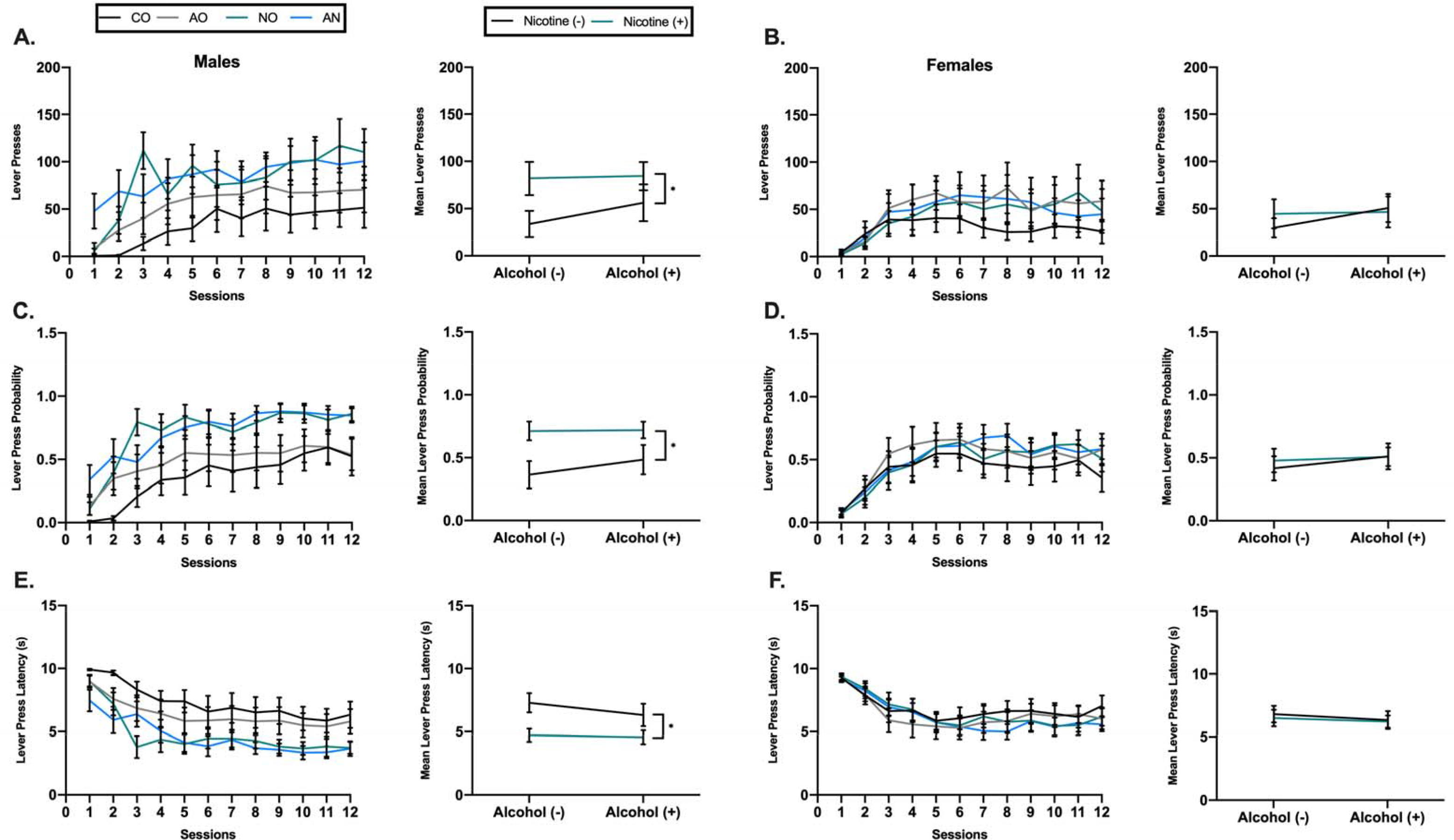
Effects of adolescent nicotine vapour and alcohol exposure on sign-tracking in adult male and female rats. Males exposed to nicotine vapour as adolescents showed A) increased lever pressing compared to CO males C) increased probability to press the lever compared to CO and AO males and E) a shorter latency to press the lever compared to CO and AO males. B, D, F) No differences were observed amongst female exposure groups in sign-tracking measures. The data is presented as mean ± SEM. * p ≤ 0.05 significant difference between males that received vapourized nicotine and males that did not receive vapourized nicotine in adolescence.

Goal-tracking was measured as food cup entries during CS+ presentation over the 12 testing sessions. Analysis of the number of food cup entries revealed a main effect of session for females [F(3.5,108.9) = 6.7; p < 0.05, η_p_^2^ = 0.177] but not males. There was no effect of nicotine vapour or alcohol exposure in either males or females **(Figure 3 A,B)**. Analysis revealed a main effect of session for the probability and latency to enter the food cup in males [F(2.4,85.6) = 10.9; p < 0.05, η_p_^2^ = 0.237], [F(2.6,91.0) = 24.2; p < 0.05, η_p_^2^ = 0.408] and females [F(2.4,74.9) = 10.6; p < 0.05, η_p_^2^ = 0.256], [F(3.1,96.2) = 25; p < 0.05, η_p_^2^ = 0.446]. An effect of nicotine vapour for males in the probability and latency to enter the food cup was observed [F(1,35) = 5.1; p = 0.03, η_p_^2^ = 0.128], [F(1,35) = 6.6; p = 0.02, η_p_^2^ = 0.158], with AN exposed males demonstrating a lower probability and longer latency to enter the food cup compared to CO (*d* = 1.1) and AO (*d* = 1.0) exposed males **(Figure 3 C,E)**. In contrast, nicotine vapour exposure had no effect in females for either the probability or latency to enter the food cup **(Figure 3 D,F).** A repeated measures ANOVA for CS- approach revealed no significant effects of session and exposure across all behavioural metrics for males and females (data not shown).

**Figure 3.**
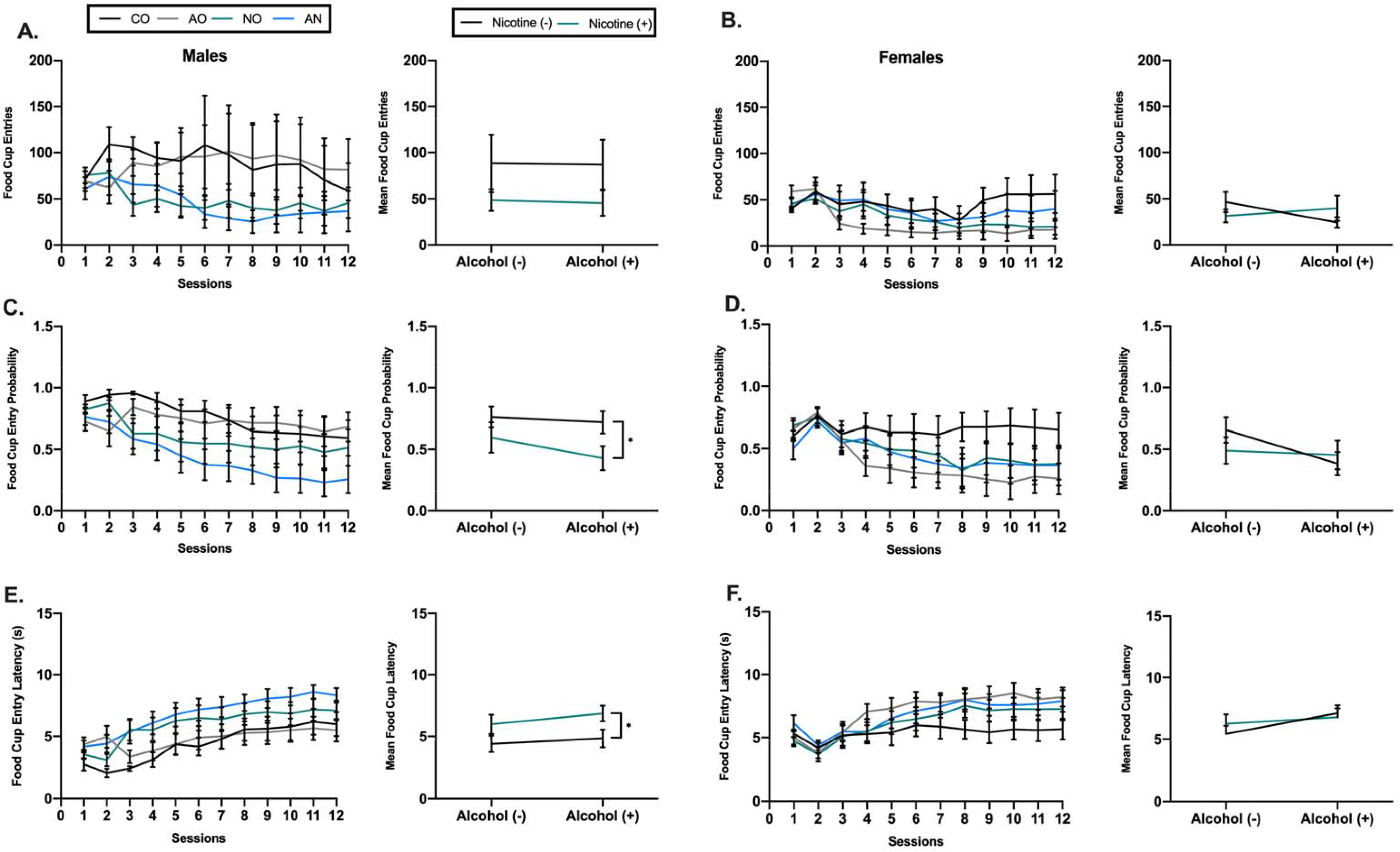
Effects of adolescent nicotine vapour and alcohol exposure on goal-tracking in adult male and female rats. A) No exposure group differences in the number of food cup entries was observed amongst males. Males exposed to nicotine vapour as adolescents showed C) decreased probability and E) an increased latency to enter the food cup, with a strong effect detected between AN males vs. CO and AO males. B, D, F) No differences were observed amongst female exposure groups in goal-tracking measures. The data is presented as mean ± SEM. * p ≤ 0.05 significant difference between males that received vapourized nicotine and males that did not receive vapourized nicotine in adolescence.

Analysis of the average PCA index score revealed a main effect of nicotine vapour [F(1,35) = 8.4; p = 0.006, η_p_^2^ = 0.194] in males exposed to vapourized nicotine. According to Cohen’s criteria, a large effect was observed in the comparison between AN and CO (*d* = 1.2) exposed males, AN and AO (*d* = 0.9) exposed males, and between NO and CO (*d* = 0.9) exposed males, where male groups exposed to vapourized nicotine had a higher PCA index score **(Figure 4A)**. No such effect was observed in females **(Figure 4B).** Co-exposure of alcohol and nicotine increased the number of male sign-trackers to 63.6% relative to male controls of 0%. Conversely, co-exposure of alcohol and nicotine had no effect on the number of female sign-trackers relative to female controls with 22.2% of co-exposed females presenting with a sign-tracking phenotype compared to 25% of female controls **(Table 1).**

**Table 1.** PCA Index Scores Data are presented as percentage of animals in each exposure group exhibiting goal tracking (GT), intermediate (Int.) or sign tracking (ST) phenotypes.

| Male |  |  |  |  | Female |  |  |  |  |
| --- | --- | --- | --- | --- | --- | --- | --- | --- | --- |
|  | CO | AO | NO | AN |  | CO | AO | NO | AN |
| **GT** | 50 | 45.4 | 22.2 | 27.3 | **GT** | 50 | 22.2 | 33.3 | 22.2 |
| **Int.** | 50 | 36.4 | 33.3 | 9.1 | **Int.** | 25 | 22.2 | 22.2 | 55.6 |
| **ST** | 0 | 18.2 | 44.5 | 63.6 | **ST** | 25 | 55.6 | 44.5 | 22.2 |

**Figure 4.**
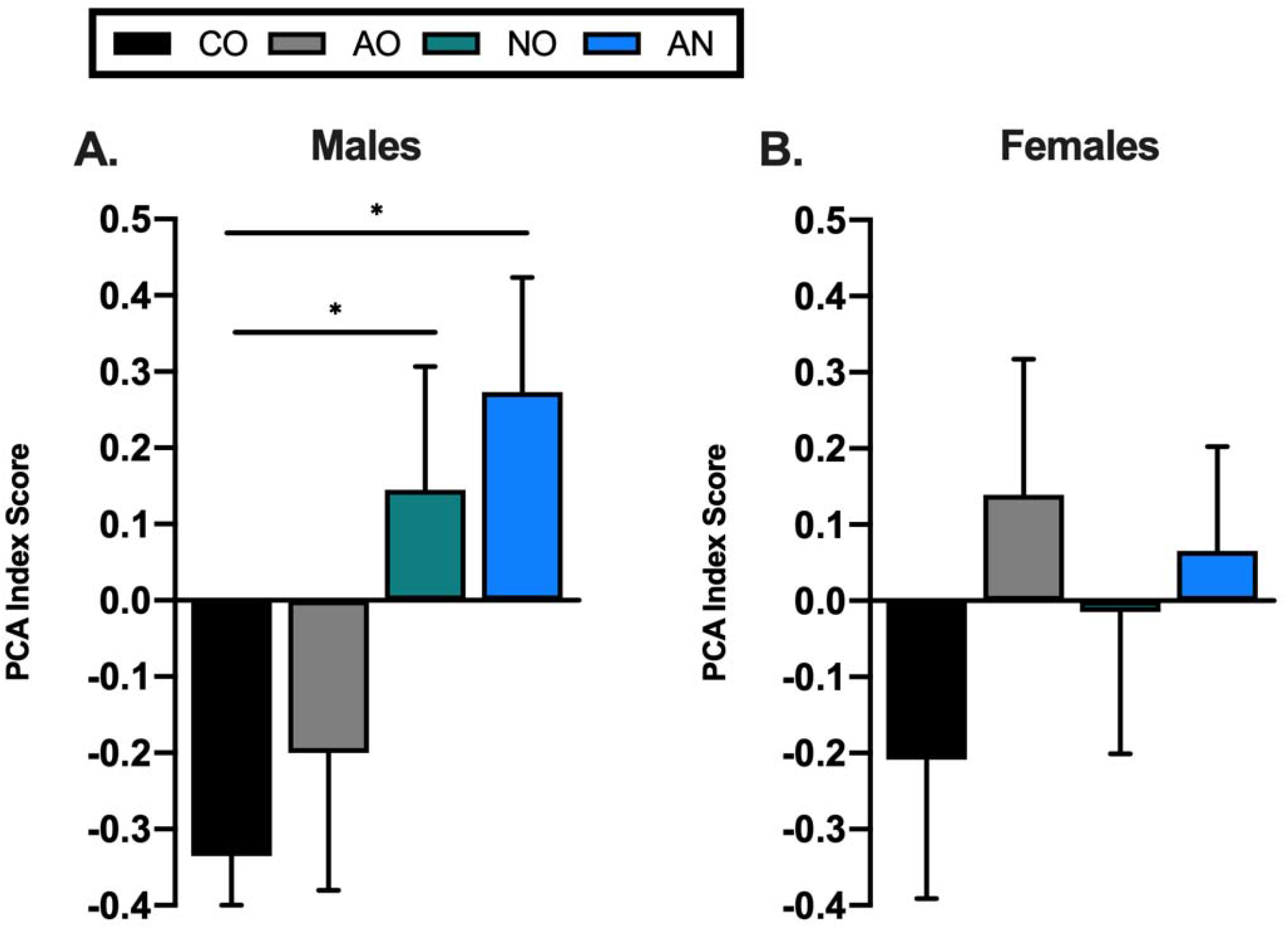
Lever press and food cup entry number, probability and latency were combined into a PCA index score for each session and averaged over the 12 PCA sessions. PCA index scores are used to classify rats as sign-trackers (STs) (score +0.3 to +1.0), intermediates (Int.) (score −0.29 to +0.29) and goal-trackers (GTs) (score −1.0 to −0.3). A) male groups exposed to vapourized nicotine in adolescence had a higher PCA index score compared to CO males. B) No significant differences for PCA index scores were detected in females. The data is presented as mean ± SEM. * p ≤ 0.05 significant difference between males that received vapourized nicotine and males that did not receive vapourized nicotine in adolescence.

### Fear Conditioning

#### Adolescent alcohol exposure in male rats impaired learning and contextual fear memory

A main effect of post-shock trial for males [F(3.6,116.4) = 26.8; p < 0.05, η_p_^2^ = 0.456] and females [F(3.1,93.4) = 22.5; p < 0.05, η_p_^2^ = 0.428] was observed on the conditioning day, indicating a progressive increase in freezing behaviour as trials proceeded. During the conditioning session, a main effect of alcohol [F(1,32) = 9.2; p = 0.005, η_p_^2^ = 0.224], nicotine vapour [F(1,32) = 4.3; p =0.46, η_p_^2^ = 0.119] and an alcohol-by-nicotine vapour interaction [F(1,32) = 8.3; p = 0.007, η_p_^2^ = 0.206] was detected in males **(Figure 5A),** where male adolescent drug exposure resulted in significant deficits in fear acquisition relative to CO males. The level of post-shock freezing in females was comparable in all exposure groups during training **(Figure 5B).** Male and female rats in each group exhibited freezing during the context test, which diminished over the course of the session. This was confirmed by a repeated measures ANOVA in which there was a main effect of epoch in males [F(4.3,138.9) = 7.1; p < 0.05, η_p_^2^ = 0.181] and females [F(3.6,106.8) = 5.7; p < 0.05, η_p_^2^ = 0.160]. Analysis revealed a main effect of alcohol in males [F(1,32) = 5.6; p = 0.02, η 2 = 0.150] where AO and AN exposed males showed less freezing behaviour and demonstrated deficits in context-related memory **(Figure 5C).** There was no main effect of exposure on context-related memory for female rats **(Figure 5D).** A main effect of trial on cue testing day was revealed for males [F(10.8,345.6) = 8.3; p < 0.05, η_p_^2^ = 0.207] and females [F(8.0,240.1) = 7.2; p < 0.05, η_p_^2^ = 0.194], indicating both sexes demonstrated extinction of the fear response over repeated tone administration. No effect of exposure on cue-related memory was detected for either male or female rats **(Figure 5 E,F).**

**Figure 5.**
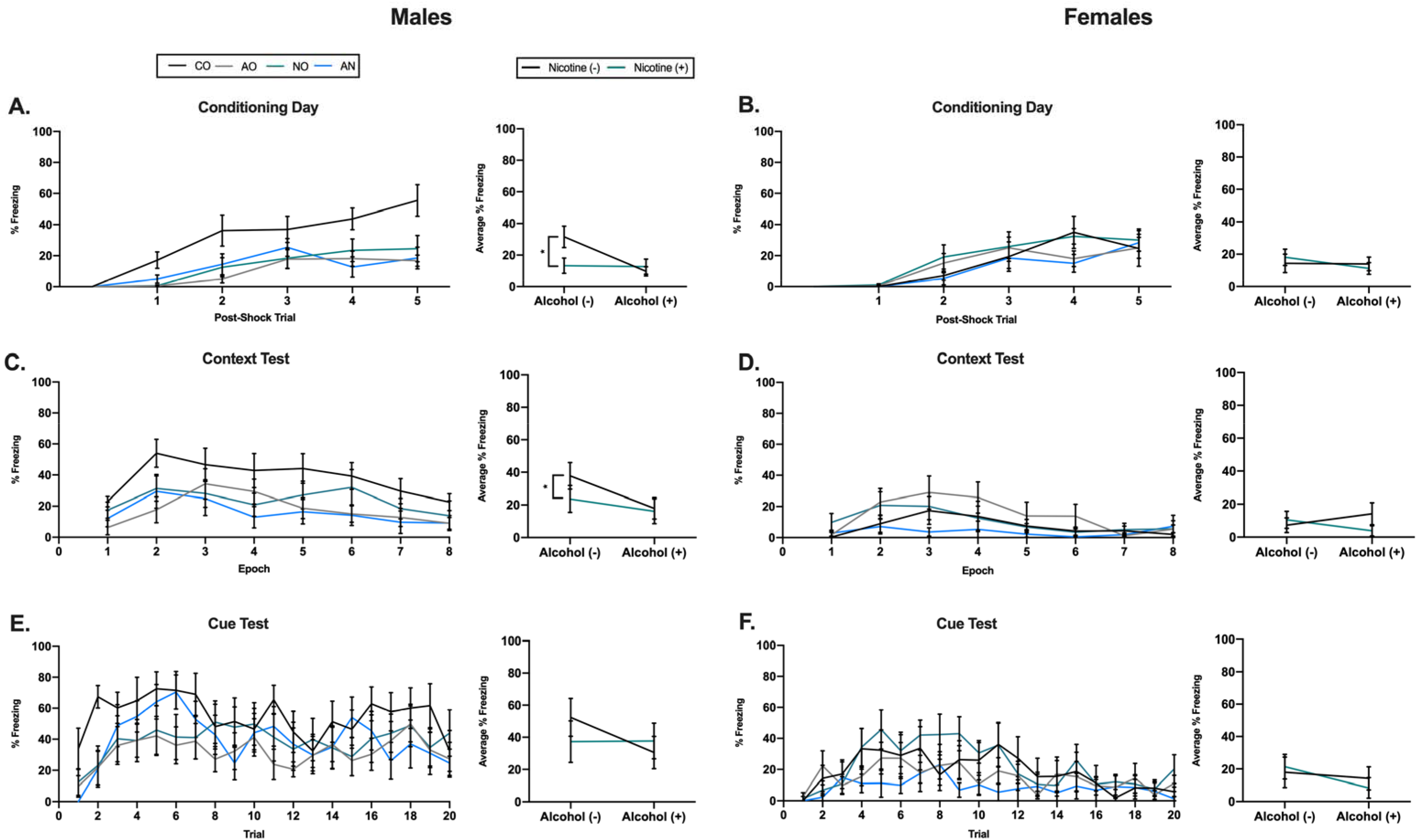
Effects of adolescent drug exposure on freezing behaviour in male and female rats. Adolescent drug exposure significantly impaired fear acquisition in A) males but not in B) females. C) Male groups exposed to alcohol showed impairments in contextual fear memory. D) No significant effects were detected in females. E, F) Adolescent drug exposure had no significant effects on freezing in males or females during the tone test session. Data is presented as mean ± SEM. * p ≤ 0.05 significant difference between A) male exposure groups and CO males and C) AO and AN males compared to CO males.

## DISCUSSION

Despite the epidemiological findings indicating frequent associations between e-cigarette use and alcohol consumption^4,5^, there is a scarcity of information regarding the long-term behavioural consequences of their co-use during adolescence. Using a preclinical model of moderate voluntary adolescent alcohol intake and passive nicotine vapour administration, the current study found that exposure to alcohol and vapourized nicotine during this critical developmental period promotes unique sex-specific effects on incentive and associative learning processes.

The present study demonstrated adult male, but not female, rats exposed to vapourized nicotine in adolescence attributed greater incentive value to reward-predicting cues, evidenced by increased approach toward the cue (i.e., CS+), and were more likely to express a sign-tracking phenotype. The current findings confirm previous observations showing that subcutaneous nicotine injections during adolescence produce long-lasting alterations in reward-associated learning in adult male rats^17^. Animals that develop the conditioned response to approach the cue are termed sign-trackers, while others that preferentially approach the reward delivery location when the cue is presented are termed goal-trackers. Several studies have shown that exposure to substances of abuse foster sign-tracking behaviours^15,16^, with sign-tracking phenotypes being often associated with behaviours such as reduced impulse control and psychomotor sensitization^32,33^. Thus, exposure to vapourized nicotine during adolescence in males appears to produce a phenotype previously associated with addiction-vulnerability. Our study is also supported by clinical findings suggesting that nicotine use increases attentional bias to drug-associated cues in smokers^34^. The ability of nicotine vapour to enhance the incentive value of reward-predicting cues may be relevant to the initiation of future smoking. A meta-analysis revealed adolescent e-cigarette users had more than three times the odds of subsequent cigarette use and four times the odds of past 30-day smoking than non-users^35^. Additionally, in a human laboratory paradigm, the exposure to passive e-cigarette use increased adolescents urge to smoke a regular cigarette^36^. Based on our observations and the aforementioned evidence, this suggests that e-cigarette use in adolescence poses as a major risk factor for future tobacco use and dependence.

Female rats appeared to be resistant to the long-lasting incentive-salience related effects observed in male rats. This contrasts previous findings that nicotine exposure in female rodents enhances the expression of sign-tracking behaviours^37^. However, these studies were performed in females injected with nicotine in adulthood, whereas repeated injection with nicotine in adolescence reduced approach behaviours in adult female rats^17^. These sex differences highlight the potential impact of route of administration, as well as the age of exposure on the long-term effects of adolescent nicotine vapour exposure on reward-related behaviours. There is also evidence that hormones influence reward-associated behaviours. In females, the rewarding effects of substances of abuse are often elevated during times of low progesterone and high estrogen^38^ and cues paired during the estrous stage have shown increased dopaminergic firing as opposed to the proestrus stage of the estrous cycle^39^. However, follow-up studies are needed to determine the mechanistic differences that underlie these sex-dependent behavioural effects.

Conversely, adolescent exposure to alcohol had no effect on sign-tracking behaviours in adulthood. This adds to the mixed literature suggesting that prior alcohol exposure can lead to either an enhanced^15,16,40^ or absent^16^ sign-tracking responses. One factor contributing to this discrepancy might be differences in task design; previously, a divergence in behavioural responses between adolescent alcohol-exposed and control animals was detected by the 14^th^ day of testing^40^, requiring more training days than our study included. Interestingly, nicotine vapour and alcohol co-exposure led to similar behavioural profiles to nicotine vapour alone in the majority of the PCA measures, suggesting that a synergistic or additive effect of nicotine and alcohol was not present for these behaviours and their combined effect was primarily driven by the actions of nicotine vapour. However, with respect to goal-tracking, males co-exposed to nicotine vapour and alcohol during adolescence, but not nicotine vapour alone, showed decreased goal-tracking behaviours relative to males exposed to alcohol alone, suggesting a potentiation of nicotine vapour’s effects by alcohol. Consistently, co-administration of alcohol and nicotine produces an additive release of dopamine in the nucleus accumbens core compared to each drug in isolation^41^, as well as behavioural disruptions not observed following alcohol administration exclusively^42^.

Male, but not female, rats exposed to alcohol, nicotine or the combination during adolescence exhibited a deficit in fear acquisition compared to controls as denoted by a reduction in the percentage of freezing time across the post-shock trials. These results are consistent with previous work suggesting that sex moderates the effects of adolescent drug exposure on cognitive function^43^. The reduced fear response in adolescent drug-exposed males may hinge on impairments in underlying learning mechanisms. Specifically, the amygdala plays a role in the acquisition of fear memory^44^, and drug use in adolescence has been documented to disrupt amygdala processes that persist into adulthood^45^, suggesting that the amygdala may be particularly vulnerable to adolescent drug exposure. The amygdala is also implicated in tone fear conditioning; however, we found adolescent drug exposure did not influence behaviour to the tone presented alone. The several nuclei that make up the amygdala each mediate different types of conditioned fear behaviour^46^. Lesions to the basolateral nuclei produce deficits in conditioned fear acquisition, while not altering auditory-cue conditioning^47^. This implies that the impairments we observed in fear acquisition, but not tone-induced freezing, might be a result of drug-induced insult to only the basolateral nuclei while sparing other nuclei necessary for tone cue-conditioning. Our results are in agreement with prior studies that found no differences in tone freezing responses in adult rats exposed to alcohol or nicotine during adolescence^20,48^. Adolescent alcohol-exposed male rats showed contextual memory deficits in adulthood, which is dependent on the hippocampus^49^. However, given that alcohol exposure compromised fear acquisition, the context memory deficits may not be due to memory retention, but a result of learning impairments. Nonetheless, these effects corroborate previous literature that have reported alcohol-induced disruptions of contextual fear conditioning during adolescence^20^.

In contrast to previous findings with nicotine^48,50^, we found no effect of adolescent nicotine vapour exposure during contextual fear conditioning in either male or female rats. However, these inconsistencies may be a result of differing study parameters such as the strain of animal, dosage used, route of administration, and dependent measure to assess fear conditioning. In fact, studies utilizing freezing as the dependent measure produced a similar lack of effects^51^, whereas lick suppression protocols showed adolescent nicotine-induced deficits^50^. These procedural differences could therefore hinder study comparisons and limit the development of hypotheses related to the effects of vaporized nicotine. As noted earlier, male rats expressed enhanced freezing behaviours compared to female rats. However, these discrepancies may not reflect genuine learning and memory deficits in females, but rather be a product of our fear assessment method. Male rats are more likely to perform inactive responses such as freezing, while females engage in active responses such as darting representing escape-like behaviour, therefore exhibiting lower levels of freezing^31^. For that reason, future studies should utilize multiple indices of fear behaviours, particularly when comparing sexes. While the exact neurobiological basis that underlies this sexual dimorphism remains unknown, these findings add to our current understanding regarding the role of sex and drug use in learning and memory.

Contrary to our predictions, we found that the exposure to nicotine vapour, alcohol or its combination in adolescence had no effect on alcohol intake or preference in adulthood. These results are in agreement with previous findings on adolescent nicotine and alcohol exposure that enforced voluntary access to alcohol in adulthood^52,53^. However, there are inconsistencies throughout the literature, where previous studies have shown early exposure to alcohol either increased^12,54^ or decreased^27^ subsequent alcohol intake. Additionally, both the exposure to nicotine and the co-exposure of nicotine and alcohol in adolescence has been reported to enhance alcohol drinking later in life^10^. These conflicting patterns of responses may be dependent on methodological distinctions such as regimen of administration during adolescence and choice of alcohol consumption paradigms used in adulthood. For instance, forced alcohol exposure has been shown to induce stress as opposed to voluntary access, and as a result increases alcohol preference^53^. Moreover, taste aversion to alcohol may account for the lack of increased alcohol consumption in the present study. Indeed, it is well established that alcohol induces an aversive effect in rodents causing the restriction of ingestion, even in those selectively bred to prefer alcohol.^55^

Consistent with previous research^54^, we found that female rats consumed significantly more alcohol than male rats. A potential explanation for this sex-specific pattern may be the fact that female rats are less sensitive to the hypnotic effects of alcohol relative to males^56^. This insensitivity would serve as a permissive feature, leading females to consume more alcohol before experiencing pharmacological feedback that would moderate their intake. Prior studies have suggested potential impacts of the female reproductive cycle and fluctuating hormones on alcohol intake with changes in the reinforcing properties of alcohol throughout the estrous cycle in rats with synchronized cycles^56^ while others have reported no impact in freely cycling rats^57^. Though we cannot be certain that our female rats are asynchronous, we saw no day-to-day variations in their drinking behaviours that would indicate a change in alcohols reinforcing effects. Thus, the present results suggest that gonadal hormone fluctuations may have minimal influence on the consumption of alcohol in our female rodents. Although animals exposed to alcohol in adolescence did not differ from controls in alcohol consumption during adulthood, within-animal differences in the consumption of alcohol during adolescence compared to later drinking demonstrated increases in preference for both male and female rats. It was also observed that female rats consumed more alcohol in adulthood compared to their intake in adolescence. Given the prominent brain development that occurs during adolescence, it is likely that alcohol sensitizes neuronal circuits involved in reinforcement (i.e. mesocorticolimbic pathway) that increases the risk of alcohol-related issues later in life^58^. Additionally, these results may suggest that adult consumption is related to solution familiarity as opposed to the effects of adolescent alcohol exposure itself. When interpreted this way, our data is in accordance with previous findings that have reported enhancement of adult alcohol consumption relative to drinking during adolescence due to solution acceptance^59^. Taken together with the literature, our findings indicate that the effect of adolescent exposure to nicotine vapour, alcohol or the combination in regulating alcohol consumption in adulthood heavily depends on multiple variables, and the complexity of these findings highlight the significance in considering intra-individual differences as opposed to solely group differences in such analyses.

The present study is the first to investigate the long-term sex-specific effects of adolescent concurrent nicotine vapour and alcohol exposure on subsequent behaviours, showing that adolescent alcohol drinking, vapourized nicotine and their combined exposure impact reward-driven and cognitive-associated behaviours later in life. We did not find additive effects of the combination of alcohol drinking and nicotine vapour; it is likely that alternate dose combinations may be needed to observe a drug interaction. Furthermore, we did not assess the impact of co-exposure on the pharmacokinetics and pharmacodynamics of individual drugs, and how they may be influenced by age or sex. This is of critical importance given the majority of our findings were sex-specific. Future studies incorporating assessments of plasma drug levels and receptor expression should be conducted to provide better insight into these observed differences. Nevertheless, our results add to the growing list of findings that highlight the sex-related variations that emerge and call attention to the importance of utilizing both sexes when measuring behavioural and cognitive outcomes. With the recent escalation of e-cigarette use among teens and its association with the consumption of alcohol, these findings underscore the importance of studying the causal consequences of e-cigarette and alcohol co-use during adolescence, and ultimately elucidate the neurobiological underpinnings that drive the effects of these drugs.

## Acknowledgments

JR, JAF, HHAT and JYK conceptualized the paper. JR, JAF, HHAT and MAT performed experiments. JR and JAF performed analyses. JR wrote the manuscript and JYK, JAF and HHAT provided manuscript revisions and finalized the manuscript for submission; all authors have given feedback on the final manuscript and approved its submission.

